# Identifying Sex Differences in Lung Adenocarcinoma Using Multi-Omics Integrative Protein Signaling Networks

**DOI:** 10.1101/2025.02.03.636354

**Authors:** Chen Chen, Enakshi Saha, Jonas Fischer, Marouen Ben Guebila, Viola Fanfani, Katherine H. Shutta, Megha Padi, Kimberly Glass, Dawn L. DeMeo, Camila M. Lopes-Ramos, John Quackenbush

## Abstract

Lung adenocarcinoma (LUAD) exhibits differences between the sexes in incidence, prognosis, and therapy, suggesting underexplored molecular mechanisms. We conducted an integrative multi-omics analysis using the Clinical Proteomic Tumor Analysis Consortium (CPTAC) and The Cancer Genome Atlas (TCGA) datasets to contrast transcriptomes and proteomes between sexes. We used TIGER to analyze TCGA-LUAD expression data and found sex-biased activity of transcription factors (TFs); we used PTM-SEA with CPTAC-LUAD proteomics data and found sex-biased kinase activity. We combined these to construct a kinase-TF signaling network and discovered druggable pathways linked to cancer-related processes. We also found significant sex biases in clinically relevant TFs and kinases, including NR3C1, AR, and AURKA. Using the PRISM drug screening database, we identified potential sex-specific drugs, such as glucocorticoid receptor agonists and aurora kinase inhibitors. Our findings emphasize the importance of considering sex and using multi-omics network methods to discover personalized cancer therapies.

## Introduction

Lung adenocarcinoma (LUAD) is the most common type of non-small cell lung cancer (NSCLC); although its prevalence in males is decreasing, it is becoming more common in younger females, and lung cancer remains the leading cause of cancer-related mortality worldwide ^1,2^. Despite advances in diagnosis and treatment, notable disparities persist, particularly in incidence and mortality rate between sexes. In 2020, adenocarcinoma was the most common type of lung cancer, accounting for 57% of cases in women and 39% in men ^3^, with males having worse prognosis ^4,5^. Treatment responses also show sex-associated variations: females often respond better to chemotherapy, whereas males typically benefit more from immune checkpoint inhibitors ^6,7^. However, the molecular mechanisms underlying these sex differences remain poorly studied and largely unclear.

Sex differences in cancer have generally been attributed to the influence of sex hormones ^8–10^. Female hormones (such as estrogens) are typically considered protective against lung cancer (despite high incidence in women), while male hormones (such as androgens) have been associated with increased risk ^11–13^. Beyond hormonal influences, genetic and metabolic factors also contribute to the comparatively better prognostic outcomes observed in females with lung cancer ^14,15^. Immune system function differences between males and females may further explain these disparities, as evidenced by sex-specific variations in the expression levels of immune-related genes ^6,16,17^. Although many studies have investigated genetic variants and gene expression in LUAD, integrating datasets spanning transcriptomes, proteomes, and post-translational modifications (PTMs) offers a unique opportunity to uncover sex-specific molecular mechanisms influencing both lung cancer risk and clinical outcomes. Ultimately, our goal was to identify potential therapeutic targets and associated agents that exhibit sex-dependent efficacy patterns.

The Cancer Genome Atlas (TCGA) is an extensive genomic repository that includes gene expression, mutation, and clinical data, crucial for LUAD research ^18^. The National Cancer Institute (NCI)’s Clinical Proteomic Tumor Analysis Consortium (CPTAC) provides complementary information including extensive genomic, proteomic, and PTM data from LUAD ^19^. Notably, CPTAC has compiled a substantial protein phosphorylation dataset, which is particularly valuable for studying kinases—key regulators of cellular signaling pathways and major targets of cancer drugs. Worldwide, over 120 small-molecule kinase inhibitors (SMKIs) have been approved for various diseases, including nearly 70 by the FDA specifically for cancer ^20^. These include targeted therapies such as EGFR inhibitors (Gefitinib, Erlotinib), PI3K inhibitors (Idelalisib, Duvelisib), and MEK inhibitors (Trametinib, Cobimetinib) ^21,22^.

In this study, we used a systems biology approach to construct sex-biased signaling networks in LUAD and subsequently identified drugs targeting key nodes within these networks that exhibit sex-biased effects in LUAD treatment (Figure 1). Specifically, we used TIGER (a TF activity estimation method) ^23^ with TCGA-LUAD to identify differentially activated TFs and then PTM-SEA (which estimates kinase activity) with CPTAC-LUAD to find sex-biased kinase activation. In the next step, we integrated the differentially activated kinases and TFs into a focused protein-protein interaction (PPI) network using the OmniPath database ^24^. The robustness of the identified sex-biased signaling network was validated using independent LUAD datasets from the Applied Proteogenomics Organizational Learning and Outcomes (APOLLO) ^25^ and Gene Expression Omnibus (GEO; GSE68465) ^26^. We then used PRISM drug screening data ^27^ to assess the therapeutic targeting potential of key nodes within these pathways, focusing on differential responses in female and male LUAD cell lines.

**Figure 1.**
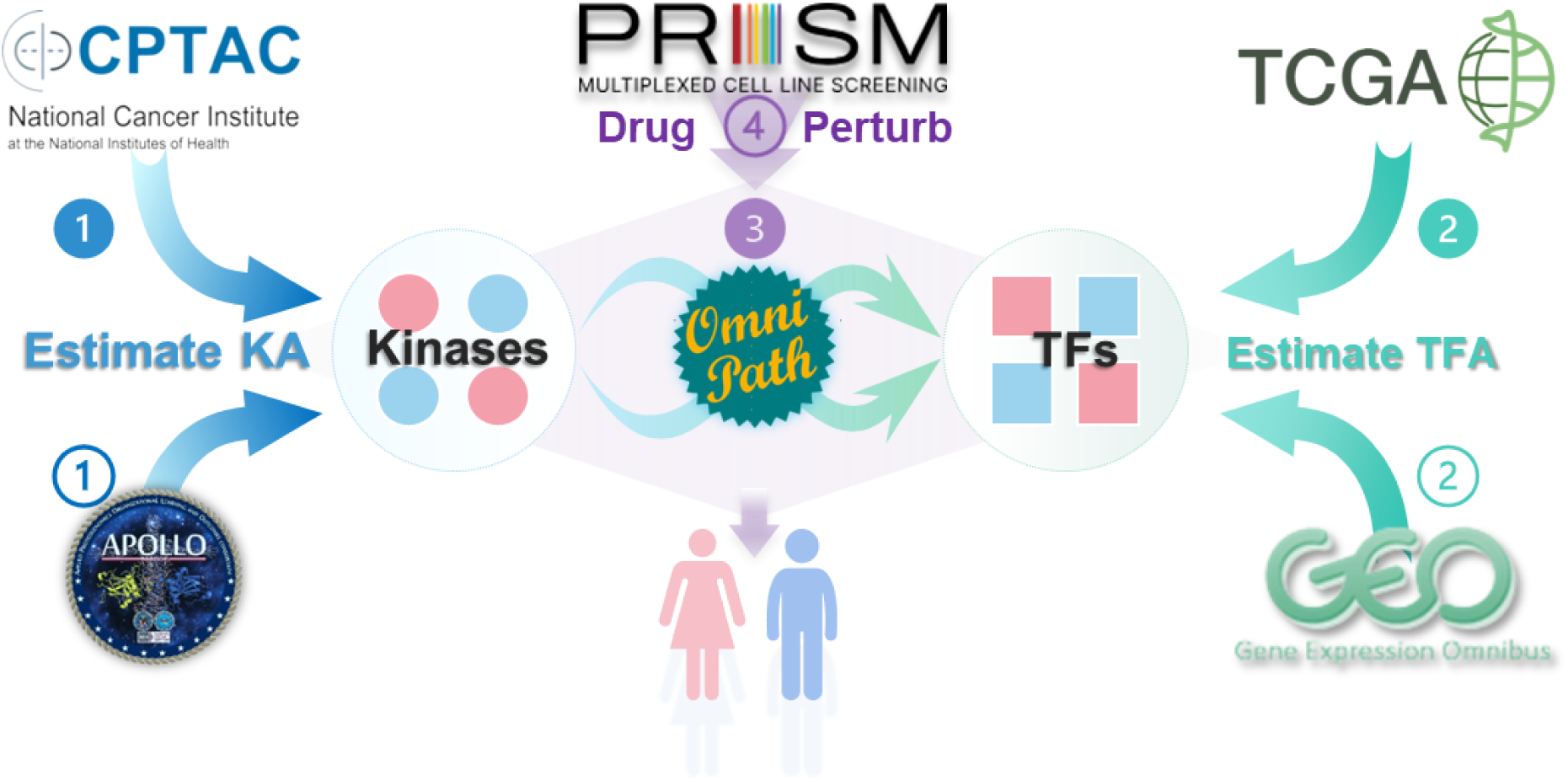
Study Overview. Illustrative flow diagram of the integrative analysis pipeline. In Step 1, protein phosphorylation data sourced from CPTAC-LUAD are used to estimate kinase activity (KA). Step 2 involves the analysis of gene expression data from TCGA-LUAD for the estimation of TF activity (TFA). Subsequently, in Step 3, differentially activated kinases and TFs are identified, with colors indicating gender specificity (red for female and blue for male). These identified kinases and TFs are then connected into a protein-protein interaction (PPI) subnetwork using the OmniPath database. The robustness of this derived subnetwork is rigorously assessed using independent validation datasets, including phosphorylation data from APOLLO-LUAD and gene expression data from GSE68465 in the GEO database. Finally, in Step 4, the PRISM drug screening dataset is employed to identify potential clinically actionable drugs, with a particular focus on developing sex-specific therapeutic strategies.

This integrative analysis allowed us to infer connections between mRNA expression and upstream protein phosphorylation, thus identifying sex-biased protein signaling pathways. Notably, the sex-biased signaling network is enriched in pathways related to cancer progression and immune response, with the latter being directly associated with poorer survival in males with LUAD. Drugs targeting key nodes in the sex-biased network, such as Aurora Kinase A (AURKA) and the Glucocorticoid Receptor (NR3C1), demonstrated significant sex-specific responses in LUAD cell lines, illustrating the potential for identifying sex-specific therapeutic strategies through multi-omics integration.

## Results

### Multi-omics landscape of sex differences in LUAD

Our multi-omics analysis of the landscape of LUAD drew upon two primary data sources. The CPTAC-LUAD dataset provided proteomic data—including protein abundance, phosphorylation, and acetylation—from 111 patients (38 females and 73 males). The TCGA-LUAD dataset had transcriptomic data from 502 patients (269 females and 233 males) after exclusion of potential outliers (Methods). The sample populations possessed diverse demographic and clinical characteristics, including sex, race, smoking status, and tumor stage (Figure 2A; Table 1). The TCGA dataset had a more balanced representation of female and male samples compared to CPTAC (Figure 2A; Supplementary Table S1). The TCGA transcriptomic dataset covered the majority of protein-coding genes, while the CPTAC proteomic dataset (Figure 2B) was a small subset of proteins encoded by TCGA transcripts. After harmonizing gene symbols to the HGNC-approved nomenclature (Methods), we found approximately 9000 genes in TCGA for which the corresponding proteins were not present in any of the other omics. The phosphorylation and acetylation datasets included an even smaller number of proteins, reflecting the fact that PTMs, which occur as chemical changes on specific amino acid residues, affect only a subset of the proteome.

**Figure 2.**
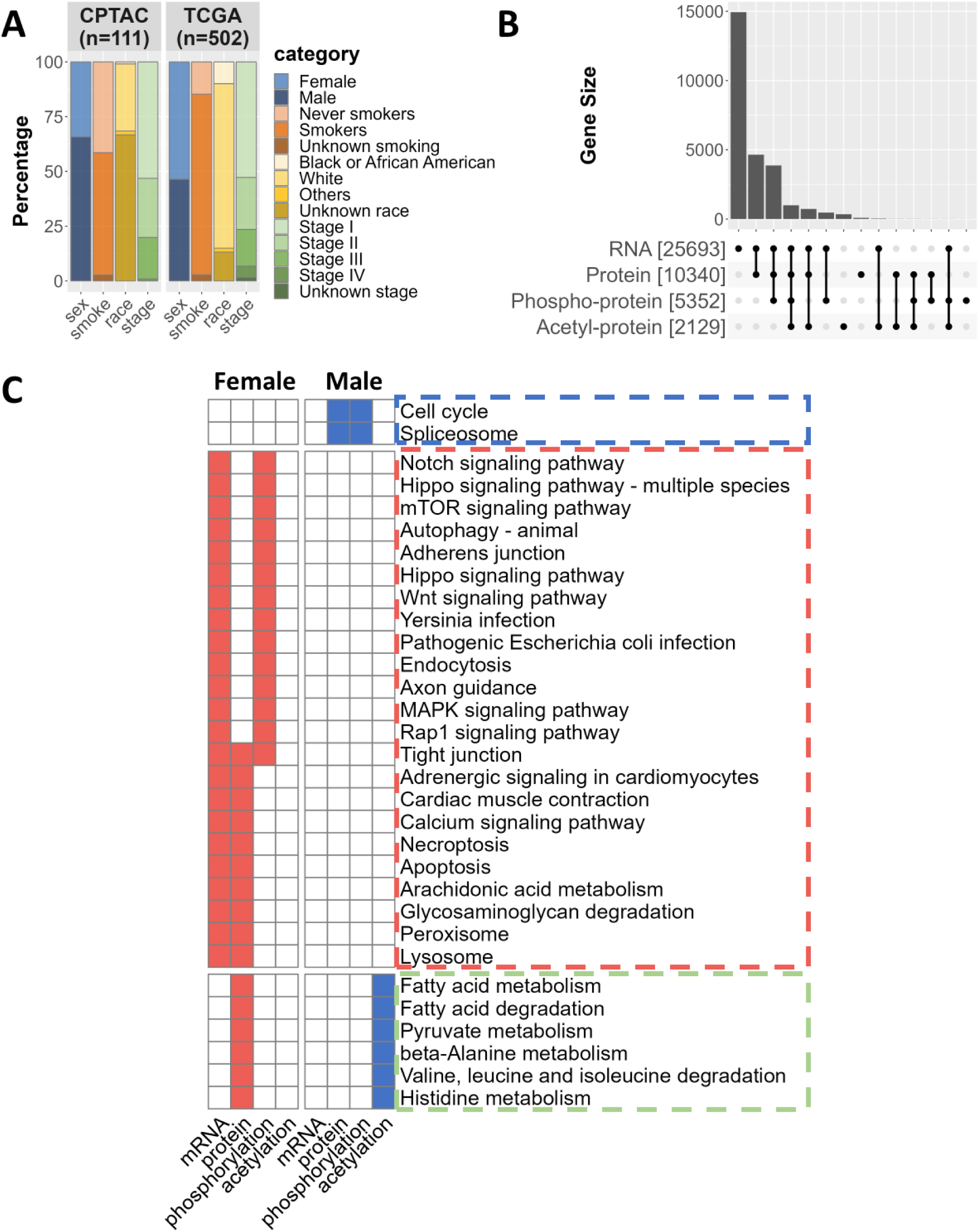
Multi-omics landscape of Sex Differences in LUAD. A. Stacked bar plot illustrating key demographic characteristics and self-reported smoking status of individuals with LUAD as documented in the CPTAC and TCGA datasets. B. UpSet plot representing the overlap of protein-coding genes represented across four omics dimensions: RNA expression, protein abundance, protein phosphorylation, and protein acetylation. All gene symbols have been aligned with the HGNC approved nomenclature (Methods). C. Visualization of the representative sex-biased KEGG pathways (adjusted p-value < 0.05, determined by GSEA or ORA) derived from multi-omics differential analysis. Pathways predominantly associated with females are marked in red, while those related to males are in blue. Pathways shared by both sexes are depicted in green.

**Table 1:** Demographic features of the discovery and validation datasets.

**Tabel 1: Demographic features of the discovery and validation datasets.**
|  | Discovery |  | Validation |  |
| --- | --- | --- | --- | --- |
|  | TCGA<br>(N=502) | CPTAC<br>(N=111) | GSE68465<br>(N=423) | APOLLO<br>(N=87) |
| **Sex** |  |  |  |  |
| Female | 269 (53.6%) | 38 (34.2%) | 214 (50.6%) | 42 (48.3%) |
| Male | 233 (46.4%) | 73 (65.8%) | 209 (49.4%) | 45 (51.7%) |
| **Age** |  |  |  |  |
| Mean (SD) | 65.5 (9.95) | 62.6 (9.61) | 64.5 (10.1) | 66.3 (10.2) |
| **Race** |  |  |  |  |
| Black or African American | 50 (10.0%) | 1 (0.9%) | 12 (2.8%) | 7 (8.0%) |
| White | 377 (75.1%) | 34 (30.6%) | 284 (67.1%) | 80 (92.0%) |
| Others | 9 (1.8%) | 2 (1.8%) | 47 (11.1%) | 0 (0%) |
| Unknown race | 66 (13.1%) | 74 (66.7%) | 80 (18.9%) | 0 (0%) |
| **Tumor stage** |  |  |  |  |
| Stage I | 265 (52.8%) | 59 (53.2%) | 145 (34.3%) | 46 (52.9%) |
| Stage II | 119 (23.7%) | 30 (27.0%) | 237 (56.0%) | 24 (27.6%) |
| Stage III | 84 (16.7%) | 21 (18.9%) | 27 (6.4%) | 15 (17.2%) |
| Stage IV | 26 (5.2%) | 1 (0.9%) | 12 (2.8%) | 0 (0%) |
| Unknown stage | 8 (1.6%) | 0 (0%) | 2 (0.5%) | 2 (2.3%) |
| **Smoking status** |  |  |  |  |
| Never smokers | 74 (14.7%) | 46 (41.4%) | 47 (11.1%) | 15 (17.2%) |
| Smokers | 414 (82.5%) | 62 (55.9%) | 286 (67.6%) | 72 (82.8%) |
| Unknown smoking | 14 (2.8%) | 3 (2.7%) | 90 (21.3%) | 0 (0%) |

We used limma ^28^ to analyze transcriptomics and proteomics data (excluding Y chromosome genes), comparing male and female samples, and adjusting for age, race, smoking status, and tumor stage; we then performed gene set enrichment analysis (GSEA) using KEGG pathway annotation to identify sex differences within these datasets (Methods). To identify robust sex-biased pathways, we intersected the results from each omics layer; however, we only observed a small number of shared pathways (Figure 2C). The modest number of shared pathways between the various data sources may be due to (1) the known weak correlation between gene expression and protein abundance ^29,30^, (2) the decreasing number of assayed genes or proteins as we move from RNA to protein abundance to protein modification, and/or (3) the limited overlap between the genes represented in different datasets (Figure 2B). Among the shared pathways found to be enriched across omics in males with LUAD were cell proliferation-related pathways, particularly cell cycle pathways. Females showed greater enrichment in cancer-related signaling pathways, including Notch, Hippo, and Wnt (Figure 2C). Metabolic pathways were enriched in female protein abundance but in male protein acetylation, consistent with acetylation’s known inhibitory effects on metabolic processes ^31–33^. A comprehensive list of the sex-biased pathways identified in each omics category is provided in Supplementary Table S2-7.

### Sex-biased protein signaling network

We used TIGER ^23^ to infer transcription factor activity in each individual in the TCGA study population. TIGER is a Bayesian matrix decomposition method that uses prior knowledge of TF-gene binding to decompose gene expression matrices, enabling the estimation of both gene regulatory networks and TF activities ^23^. We also adapted TIGER to estimate kinase activity, replacing the input gene expression matrix with a protein phosphorylation matrix and substituting the TF-gene binding prior with a kinase-substrate binding prior from the OmniPath database ^24^. Simultaneously, we used the PTM-SEA ^34^ algorithm to estimate kinase activity. PTM-SEA is a modified version of the GSEA algorithm designed to perform site-specific signature analysis ^35^. It uses the PTM signatures database (PTMsigDB) to score PTM site-specific signatures, such as those for protein phosphorylation, directly from a protein phosphorylation matrix ^34^. The use of two different algorithms (TIGER and PTM-SEA) to analyze the same dataset (phosphorylation) enabled us to combine the two results to get a robust kinase activity estimation (Methods).

In our TF activity analysis, NFKB1 and NR3C1 emerged as key TF drivers in female and male patients, respectively (Figure 3A). NFKB1, also known as nuclear factor kappa-light-chain-enhancer of activated B cells, is crucial in regulating the immune response to infection ^36^. NR3C1, the glucocorticoid receptor, mediates glucocorticoids’ effects, significantly influencing inflammation and immune responses ^37^. In using TIGER and PTM-SEA for kinase activity estimation, we identified two pivotal kinases, AURKA and MAPK14, as playing vital roles in female and male patients, respectively (Figure 3B). AURKA (Aurora kinase A) is a serine/threonine kinase critical for mitosis and cellular proliferation, frequently dysregulated in cancers and contributing to their progression ^38^. MAPK14 (p38 alpha) is a key MAPK family member involved in tumor biology, regulating survival, proliferation, metastasis, and therapy response, as well as stress and inflammation signaling ^39^.

**Figure 3.**
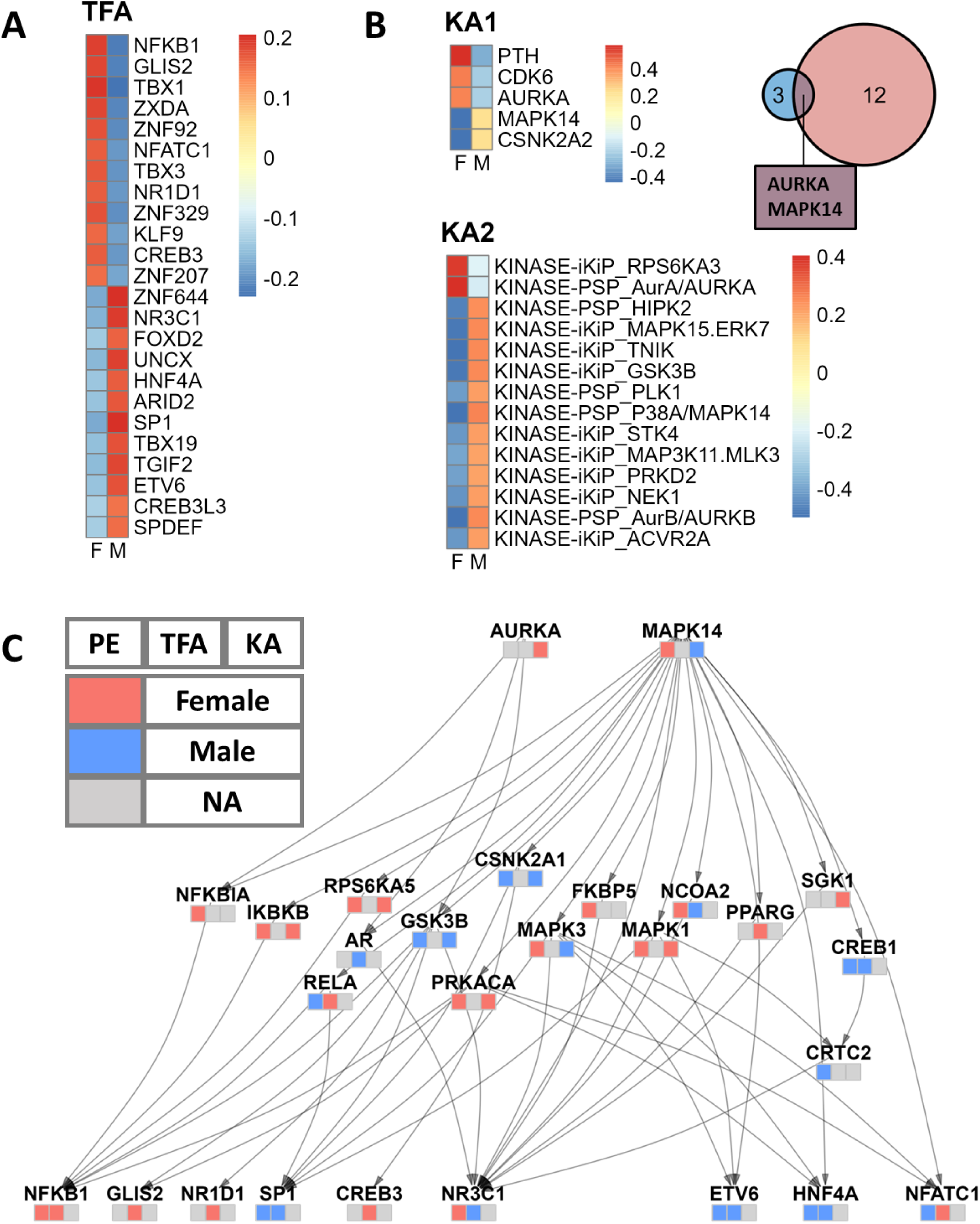
Sex-Biased Protein Signaling Network. A. Sex differences in TF activity, analyzed using the TIGER method. TFs of interest were selected based on the limma differential analysis with an adjusted p-value less than 0.05. Color indicates the average TF activity of female and male patients. B. Sex differences in kinase activity were identified using TIGER (KA1) and PTM-SEA (KA2) analyses. KA1 identified 5 kinases with an adjusted p-value cutoff of <0.25, while KA2 identified 14 kinases with an adjusted p-value cutoff of <0.1. The Venn diagram illustrates the intersection of kinases from both analyses. Heatmap color indicates the average kinase activity of female and male patients. C. The sex-biased protein signaling network, constructed using the OmniPath PPI network database, links the kinases of interest to the TFs of interest. Each node is annotated with a heatmap indicating the direction of change in protein expression (left), TF activity (middle), and kinase activity (right). A red tile indicates higher levels in females, a blue tile indicates higher levels in males, and a gray tile indicates that the gene information is missing in this channel. PE: Protein Expression; TFA: Transcription Factor Activity; KA: Kinase activity.

To integrate TF activity results from TCGA with kinase activity results from CPTAC, we curated a focused signaling network using the OmniPath protein interaction database ^24^ (Figure 3C). The network, constructed with the OmniPathR package (version 3.8.0), connects the two sex-biased kinases we identified (AURKA and MAPK) to all the sex-biased TFs. To balance comprehensiveness with sparsity and interpretability, we included paths up to three steps in length. Notably, the network highlighted the androgen receptor (AR) as a significant sex-biased intermediate node, consistent with the role of steroid sex hormones in LUAD. Over-representation analysis (ORA) on the network nodes revealed significant over-representation of cancer-related KEGG signaling pathways, such as MAPK, Wnt, mTOR signaling, and PD-1 checkpoint, along with distinctly sex-biased pathways like the estrogen signaling pathway (Supplementary Figure S1; Supplementary Table S8).

It is worth noting that when labeling the nodes by their protein expression and TF/kinase activity, we observed instances where information was missing or conflicting, emphasizing the complexity of biological systems and the necessity of a multi-omics approach. For instance, while AURKA’s protein expression is below the detection threshold and therefore missing, our method effectively identified it as displaying female-biased kinase activity.

### Sex-biased survival outcomes are associated with immune response

To investigate whether sex differences in protein signaling networks might explain the better survival outcomes observed in females with LUAD, we performed survival analysis using sex-biased signaling proteins (those in Figure 3C) and patient survival data from CPTAC. We began with an over-representation analysis of the previously identified sex-biased signaling proteins using the Gene Ontology Biological Processes (GO-BP) database, revealing 22 significantly enriched GO terms (adjusted p-value < 0.05; Supplementary Table S9). For each GO term, we computed an overall GO term score as the average protein abundance of the associated proteins. We then fit a Cox PH model for each GO term score, including sex and GO term scores as the main effects and testing their interaction term, while adjusting for age, race, smoking status, and tumor stage (Methods). Our analysis revealed that the top five most significantly sex-associated GO term scores (sex*score interaction term) are all immune-related, including scores for “regulation of defense response” (p-value = 0.02) and “inflammatory response” (p-value = 0.04). These scores were associated with better survival outcomes in females but showed little to no significant impact in males (Supplementary Figure S2).

A limitation of the survival analysis using the CPTAC data is the absence of certain protein abundance measurements and incomplete treatment information. To partially address this, we conducted an additional survival analysis using TCGA data, focusing on the target genes of our protein signaling network by constructing a kinase-TF-gene tripartite network. We first selected the top 200 target genes from the nine transcription factors shown in Figure 3C, based on their node indegrees (the sum of TIGER absolute edge strengths). We then repeated the GO term over-representation and Cox PH model analyses for each of the significant GO terms (Supplementary Table S10). Consistent with the results on CPTAC, the most significant GO terms associated with sex differences in survival outcomes (sex*score interaction term; p-value < 0.05) were immune-related (Supplementary Figure S3). The concordant findings between the CPTAC analysis (examining upstream signaling proteins) and TCGA analysis (examining downstream target genes) indicate a critical role for immune responses in driving sex-biased survival outcomes and emphasize the strength of our multi-omics analysis.

### Sex-biased regulation of histone acetylation

In cancer, aberrant histone acetylation can lead to the inactivation of tumor suppressors or the activation of oncogenes ^40–42^. Histone acetyltransferases (HATs) and histone deacetylases (HDACs), the key regulators of histone acetylation, have also been implicated in driving sex differences in both normal tissues and cancers ^43–45^. Recently, Saha and colleagues reported that Panobinostat, an HDAC inhibitor, may exhibit greater efficacy in males with LUAD because in their gene regulatory network models, its target, *CDKN1A*, is estimated to be under weaker regulatory control in males and thus more easily perturbed ^15^. Nevertheless, the differential roles of HATs and HDACs in regulating histone acetylation and altering downstream transcription between males and females with LUAD remain poorly understood.

CPTAC provides detailed information on histone acetylation, offering an unparalleled opportunity to investigate the sex-biased, histone-associated regulatory effects in LUAD. We used Least Absolute Shrinkage and Selection Operator (LASSO) regression with the CPTAC data to infer a network involving HATs and HDACs and site-specific histone acetylation levels. Specifically, we treated each histone acetylation site as the dependent variable and the set of HAT and HDAC proteins as a multivariate set of predictors, then applied LASSO regression with bootstrap to fit a penalized linear model (Methods).

We observed distinct sex-biased patterns of histone acetylation characterized by numerous positive relationships (positive LASSO coefficients) between the abundance of HATs and various acetylation sites (Figure 4). The *EP300* gene encodes p300, an HAT that plays a role in regulating cell proliferation and differentiation. We found that *EP300* has a positive relationship with three specific histone acetylation sites exclusively in the female group. This is consistent with reports of sex-biased *EP300* activity in LUAD ^46^, suggesting that *EP300* may play a role in female-specific epigenetic regulation. In contrast to the female-specific role for *EP300*, *HAT1* and *NCOA1* have a male-specific positive relationship with histone acetylation sites, indicating they may play a role in determining sex-biased disease processes in individuals with LUAD.

**Figure 4.**
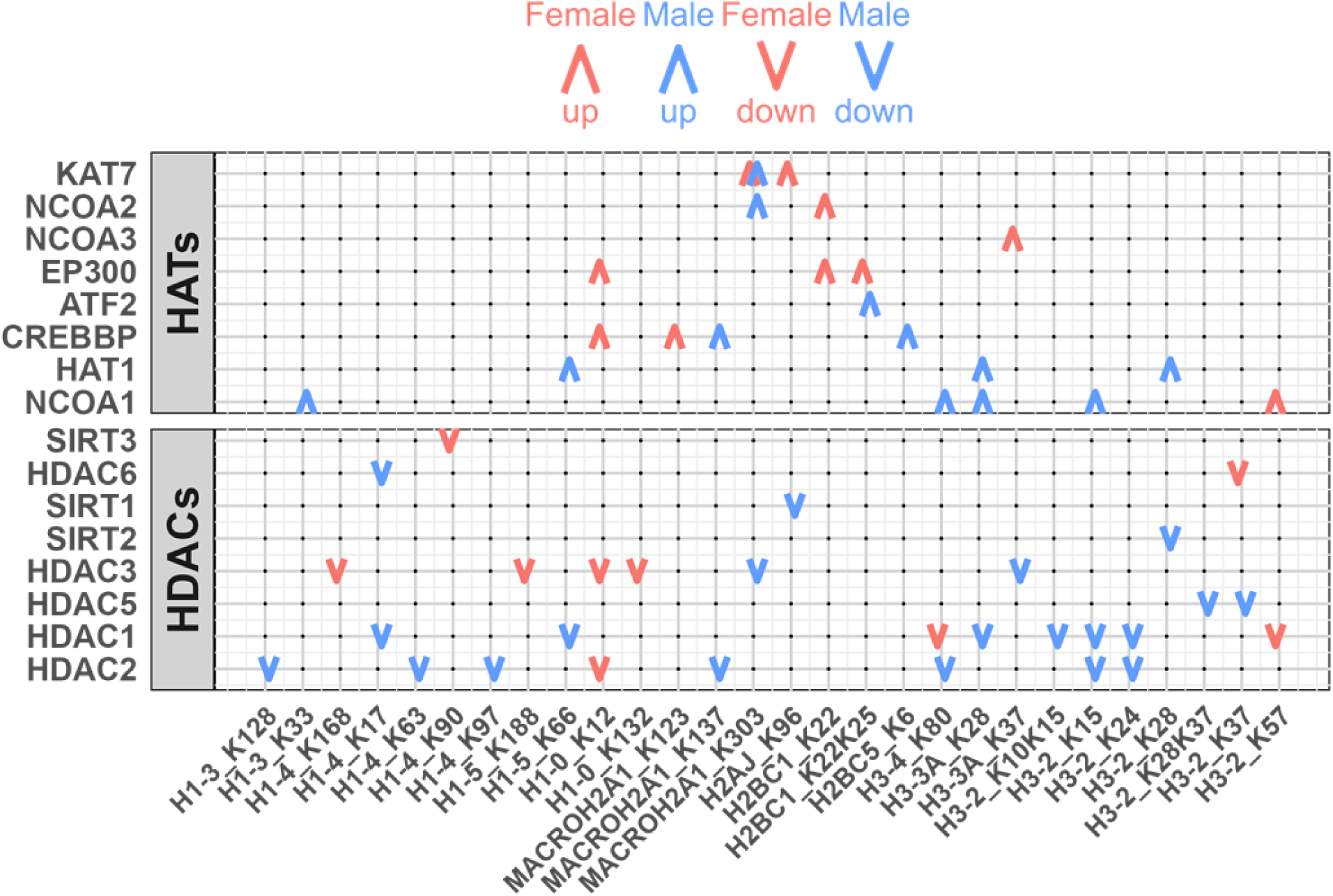
Sex-Biased Regulation of Histone Acetylation. Regulation of histone acetylation sites by HATs and HDACs. Red and blue colors represent females and males. Upward and downward arrows represent positive and negative conditional associations between regulators and histone acetyl sites. Lasso regression with bootstrap was used to select significant associations (Methods).

HDACs, known for their role in reducing acetylation levels ^47^, also showed sex-biased patterns. As expected, in both sexes we found negative relationships (those with negative LASSO coefficients) between HDAC abundance and histone acetylation levels (Figure 4). However, the magnitude of these negative relationships was more pronounced in the male group (Supplementary Figure S4; Wilcoxon signed rank test, p-value < 0.05), suggesting increased HDAC-mediated deacetylation activity in males. This finding aligns with previous reports of greater efficacy of HDAC inhibitors in males ^15^ and underscores the need for further functional and mechanistic investigations into HDACs to define their mechanistic role in LUAD development and, as described below, to understand whether HDAC inhibitors might exhibit sex biases in therapeutic effectiveness.

### Sex-biased molecular signatures of clinically actionable proteins

The signaling network model we deduced (Figure 3C) captures sex-biased patterns with key nodes that suggest potential sex-specific therapeutic strategies. We used the PRISM drug screening database ^27^ to search for small molecule drugs that might have differential inhibitory effects on male and female LUAD cell lines. Using the Wilcoxon Rank Sum Test, we identified drugs targeting nodes in our signaling network with sex-biased small molecule responses.

Specifically, sixteen drugs targeting seven proteins demonstrated statistically significant inhibition of either male or female LUAD cell lines (p-value < 0.05; Figure 5A). We did not adjust p-values in this exploratory analysis, as our goal was to identify a broad range of candidate drugs with potential sex-biased effects. The actual efficacy of these drugs should be rigorously validated through well-designed experimental studies.

**Figure 5.**
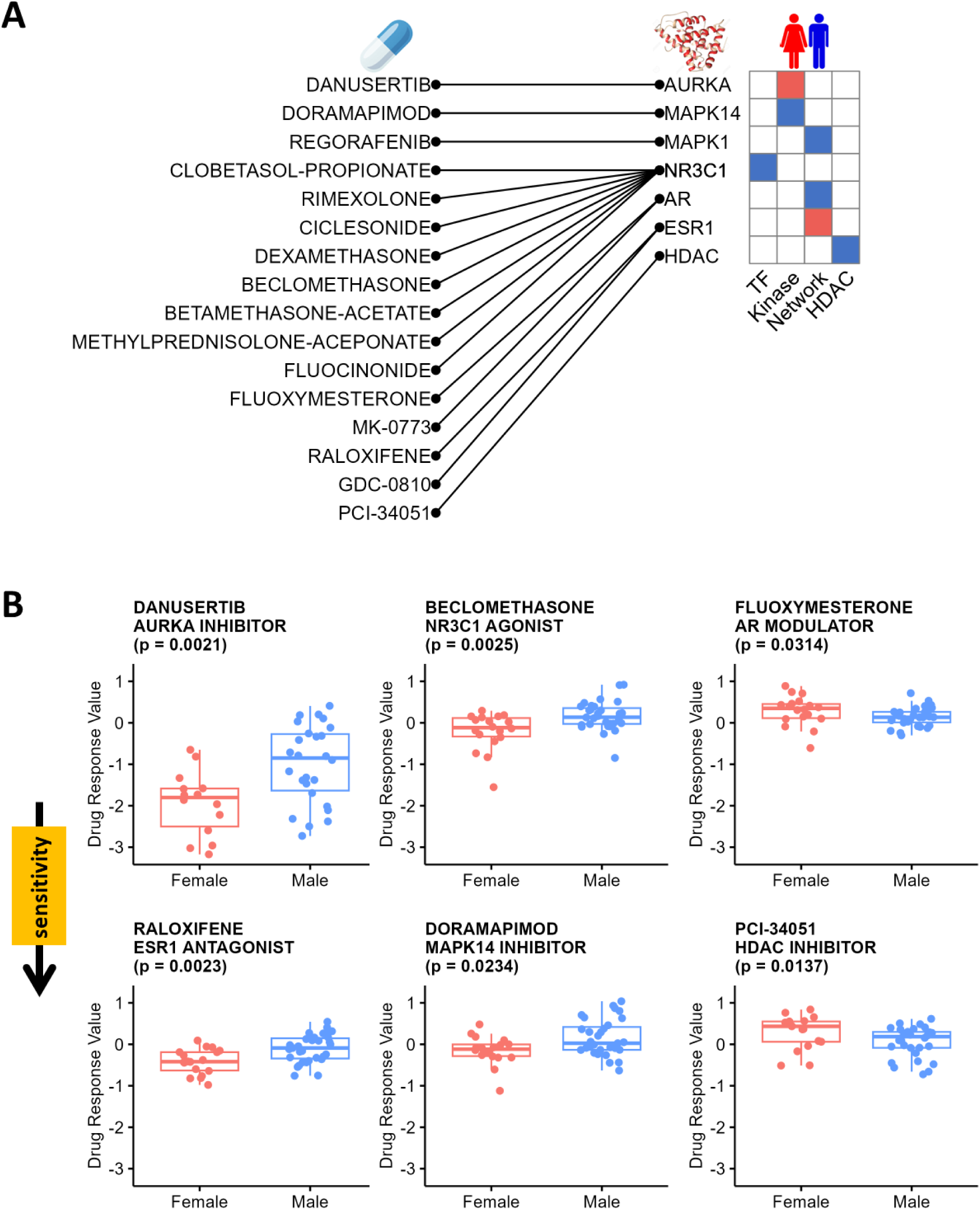
Sex-Biased Molecular Signatures of Clinically Actionable Proteins. A. The mapping of sex-biased small-molecule drugs to their associated clinically actionable proteins (left), alongside the observed sex biases in these clinically actionable proteins across various analyses (right). TFs and kinases are identified through activity analysis; network analysis uncovers intermediate proteins connecting kinases to TFs; HDAC analysis highlights sex-biased HDACs. B. Validation of sex-biased therapies using the PRISM drug screening database. Boxplots show drug response values for male and female LUAD cell lines treated with Danusertib, Beclomethasone, Fluoxymesterone, Raloxifene, Doramapimod, and PCI-34051, analyzed using Wilcoxon Rank Sum Test.

One of the most significant drugs was Danusertib (p-value = 0.0021; Figure 5B), an aurora kinase inhibitor that showed greater inhibitory effects in female LUAD cell lines than in male cell lines, consistent with higher AURKA activities in females. Treatment with NR3C1 agonists, including Beclomethasone (p-value = 0.0025; Figure 5B) and seven other synthetic corticosteroids (Supplementary Figure S5), exhibited higher sensitivity for female cell lines, consistent with our models’ conclusions that males have stronger NR3C1 activity. We also found that sex steroid hormone receptors, including AR and ESR1, were differentially targeted by several modulators, agonists, and destabilizers (p-values < 0.05; Figure 5B; Supplementary Figure S6). MAPK14 and MAPK1 inhibitors also have a sex-biased effect (p-values < 0.05; Figure 5B; Supplementary Figure S7). These drug screening results are all consistent with the sex-biased TF and kinase activities identified through our signaling network analysis.

Because we observed sex differences in relationships between HDACs and histone acetylation sites, we also tested HDAC inhibitors for sex-biased responses and identified one compound, PCI-34051, exhibiting a statistically significant higher sensitivity treatment effect in male LUAD cell lines compared to females (p-value = 0.0137; Figure 5B).

As a methodological note, we emphasize that these compounds were all found through our integrative network and histone acetylation analyses. Due to the limited throughput of proteomic data, identifying these proteins via simple differential expression proved challenging (Supplementary Figure S8). This underscores the clinical relevance of sex-biased molecular signatures and the efficacy of systems biology approaches in addressing complex biological questions.

### Independent validation of our results

We analyzed LUAD gene expression from GSE68465 (214 females and 209 males) and protein phosphorylation data from the APOLLO-LUAD project (42 females and 45 males); sample numbers reflect those remaining after filtering outliers (Methods). However, in analyzing differential gene expression, protein abundance, and protein phosphorylation, we found discrepancies between our discovery and validation datasets (Supplementary Figure S9). For instance, the Spearman correlation of limma’s t-statistics between CPTAC-LUAD and APOLLO-LUAD for the protein phosphorylation was notably low, a pattern that was also evident in the differential protein and gene expression levels (Spearman correlations = -0.086, -0,017, and - 0.056, respectively; Supplementary Figure S9A). This low concordance between CPTAC-LUAD and APOLLO-LUAD complicates the validation of driver proteins but may be due to the relatively small sample sizes in APOLLO and the imbalance between males and females in CPTAC. Indeed, with larger sample sizes, we saw better correlation between differential gene expression in TCGA-LUAD and GSE68465 (Supplementary Figure S9B; Spearman correlation = 0.34).

Although the above explorative analysis showed consistent discrepancies between CPTAC and APOLLO, we reconstructed the signaling network using the validation dataset (APOLLO and GSE68465) to assess whether the mechanisms observed in the discovery dataset (CPTAC and TCGA) hold true. Indeed, key regulators, including AURKA, NR3C1, and AR, exhibited sex-biased activities in the validation dataset (p-values = 0.03, 0.07, and 0.02, respectively; Supplementary Figure S10), supporting some of the findings in the discovery phase. At the same time, discrepancies between datasets remained (Supplementary Figure S9-10). Nonetheless, TIGER integrates multi-gene signatures representing known TF or kinase binding events, allowing it to infer robust activity levels despite dataset inconsistency. Our validation results further reinforce the role of AURKA, NR3C1, and AR as pivotal sex-biased network elements and also provide support for the PRISM drug findings reported above.

## Discussion

In LUAD, there are well-known sex differences in disease risk, progression, and therapeutic outcomes. Although this bias is certainly multi-factorial and includes genetics, hormone levels, environmental exposures, and other factors, the precise molecular mechanisms have remained elusive ^48^. By analyzing differential gene expression, protein abundance, phosphorylation, and acetylation across all chromosomes excluding the Y, we identified a handful of critical sex-biased signaling and metabolic pathways that likely also contribute to the differences between the sexes and can be more effectively targeted therapeutically based on sex.

Although there have been several omics analyses of sex bias in LUAD ^49–51^, these have generally found few differences in gene expression by sex and rarely shed light on sex biases in risk or outcomes. In contrast, studies that have analyzed changes in TF-gene regulatory networks have discovered meaningful differences not found using other methods including differential expression or coexpression analysis ^15,16,52–54^—and notably in the analysis of sex differences in LUAD ^15^. In particular, the LUAD study inferred gene regulatory network models for each individual and then compared networks of biological males and females in both healthy lung tissue and LUAD samples, identifying differential transcriptional targeting of genes.

Here, we took a complementary approach to analyzing sex differences in LUAD by focusing on integrating information that included CPTAC proteomics and TCGA transcriptomics data, to infer signaling networks that exhibit sex-biased patterns. The sex-biased signaling network we inferred was enriched for proteins associated with cancer progression and immune response, consistent with the findings reported by Saha and colleagues in analyzing TF-gene regulatory networks ^15^. Not surprisingly, we found that immune response is associated with better survival outcomes in females. Delving more deeply into the network, we were able to identify potential therapeutic targets represented as key nodes in the inferred signaling network. Using the PRISM database, we found sixteen drugs that differ in efficacy when used with cell lines derived from tumors in males and females.

Among these key nodes for which we found therapeutic candidates, many are known to play a sex-biased role in other cancers. The expression of AURKA has been reported to be higher in females with glioblastoma ^55^. This is somewhat surprising because AURKA is a proliferation marker, but females with glioblastomas generally have better outcomes than their male counterparts ^56^. Nevertheless, Danusertib is a potent inhibitor of aurora kinases with some evidence of efficacy in cancer treatment ^57,58^. Our network-seeded PRISM search identified Danusertib as a sex-biased therapeutic, with higher inhibition in female cell lines than in male cell lines.

NR3C1 encodes the glucocorticoid receptor and NR3C1 knockdowns have been shown to inhibit the proliferation and migration of clear cell renal cell carcinoma in laboratory and mouse model studies ^59^, although no sex biases have been reported. However, NR3C1 deficiency has been linked to sex-dependent DNA methylation changes in murine models ^60^, admitting a possible sex-dependent role here as well. Our analysis indicates a sex bias in the therapeutic efficacy of NR3C1 agonists, such as Beclomethasone, consistent with our observation of higher estimated NR3C1 activity in males.

The androgen receptor, AR, almost certainly contributes to sex biases; androgens have been implicated in sex bias in a number of cancers through interactions with the CD8+ T cell exhaustion program ^61^. Our PRISM analysis also identified several small molecules exhibiting sex-differential targeting of AR.

HDACs regulate gene expression by deacetylating histones, leading to chromatin condensation. We found evidence that suggests sex differences in HDAC activity, which may alter chromatin accessibility and contribute to sex-biased hormonal or other regulatory processes in disease ^43,62–64^. Further, the Y-encoded protein, KDM5D, interacts with the Sin3–HDAC complex, contributing to male-biased cancer progression ^44^. Both the analysis presented here and prior work by our group ^15^ underscore the potential importance of HDAC and its inhibitors as sex-specific therapeutics.

This study does have some limitations. First, the proteomics data are sparse, with many missing measurements across the proteome, including many proteins that are known to play a role in cancer. This missingness restricts our ability to fully investigate sex-biased protein signaling through phosphorylation or acetylation at the proteome-wide level. Second, our network integration method used the OmniPath PPI network, but a LUAD-specific (or an individual-specific) PPI network accounting for genetic variants and protein isoforms ^65^ might lead to greater insight. Third, although adjustments were made for clinical and demographic covariates like age, race, smoking history, and tumor stage, our analysis may still be influenced by other factors such as cellular and genetic heterogeneity, or unobserved clinical phenotypes and risk factors, including hormonal effects, lifestyle habits, and environmental exposures. Despite these limitations, our well-designed multi-omics discovery approach and independent validation strategy together with the biological relevance of our findings provide a level of confidence in the results we report here, particularly the potential for sex-specific therapeutic options.

In summary, this study underscores the importance both of carrying out integrative, multi-omics analyses and of considering biological sex in analyzing cancer processes. For the former, combining gene regulatory network and protein signaling network analyses appears to be a particularly effective strategy for identifying potential cancer drug candidates, complementing the deep functional insights provided by gene regulatory network analysis. For the latter, addressing the longstanding failure to consider differences between the sexes in disease mechanism and therapeutic strategies is clearly essential if we are to continue to make progress in improving cancer outcomes for all individuals.

## Methods

### Data download and preprocessing

We downloaded the TCGA-LUAD gene expression data and clinical data using the “recount3” R package (version 1.10.2) ^66,67^. We extracted TPM normalized gene expression data using the “getTPM” function in the “recount” R package (version 1.26.0) ^68^. TPM values were then log_2_ transformed to expression scores. We excluded lowly expressed genes using the “filterByExpr” function in the “edgeR” R package (version 3.42.2) ^69^ using default parameters. We then removed recurrent tumor samples and samples from adjacent normal tissues, keeping only primary tumor samples. For individuals with multiple samples, we kept only the sample with the greatest sequencing depth. Finally, we removed twelve outlier samples that appeared to exhibit batch-dependent effects based on PCA. Additionally, using PCA on Y chromosome genes, we identified and eliminated two misannotated female samples. The subsequent analyses used the expression data on 25693 genes in 502 samples (269 females and 233 males).

CPTAC-LUAD ^19^ protein abundance, phosphorylation, acetylation data, and clinical data were downloaded using the “cptac” Python package (version 1.5.1) ^70^. We removed adjacent normal samples, keeping only primary tumor samples. We filtered the data to eliminate samples that had greater than 50% missing values for each feature (abundance, phospho-site, acetyl-sites); for samples with fewer than 50% missing values, missing values were imputed using the “impute.knn” function from the “impute” R package (version 1.74.1) ^71^. This process left 10340 proteins, 20773 phospho-sites, and 5899 acetyl-sites in 111 samples (38 females and 73 males) that were used for the subsequent analyses.

We downloaded GSE68465 ^26^ LUAD gene expression data using the R “GEOquery” package (version 2.62.2) ^72^. This dataset contained LUAD specimens from the University of Michigan Cancer Center (178 samples), Moffitt Cancer Center (79 samples), Memorial Sloan-Kettering Cancer Center (104 samples), and the Dana-Farber Cancer Institute (82 samples). Principal component analysis on the gene expression data demonstrated distinct clusters corresponding to these sample sources, thus exhibiting a strong batch effect; expression data was batch-corrected using the “ComBat” function implemented in the “sva” R package (version 3.48.0) ^73^. We also removed six samples that were mis-annotated as “female,” and 14 samples mis-annotated as “male” based on PCA of Y chromosome gene expression. After cleaning, 13179 genes in 423 samples (214 females and 209 males) that were used for subsequent analyses.

APOLLO-LUAD ^25^ Level3 phosphorylation data and clinical data were downloaded from the GDC Data Portal for the APOLLO project (https://gdc.cancer.gov/about-data/publications/APOLLO-LUAD-2022). We used the official pre-processed and imputed phosphorylation dataset, comprising 2138 phospho-sites across 87 samples (42 females and 45 males), for subsequent analyses.

The PRISM ^27^ drug response data “Repurposing_Public_23Q2_Extended_Primary_Data_Matrix”, drug information data “Repurposing_Public_23Q2_Extended_Primary_Compound_List”, and the cell line information data “Repurposing_Public_23Q2_Cell_Line_Meta_Data.csv(23Q2)” were downloaded from the depmap portal (https://depmap.org/portal/download/all); we limited our analysis to LUAD cell lines by setting “DepmapModelType” to “LUAD”.

HGNC gene symbols were downloaded from https://ftp.ebi.ac.uk/pub/databases/genenames/out_of_date_hgnc/tsv/hgnc_complete_set.txt. All gene and protein names were mapped to the HGNC symbols.

### Differential expression analysis

We performed differential feature level analyses for mRNA, protein, phosphorylation, acetylation, as well as kinase and TF activities, using the limma package (version 3.56.1) ^28^. In these analyses, Y chromosome genes were removed, and age, race, smoking status, and tumor stage were adjusted as covariates. The adjusted p-value was calculated using the Benjamini-Hochberg procedure ^74^ to control for multiple testing.

### Pathway enrichment analysis

We performed KEGG pathway ^75^ analysis to assess RNA- and protein-level enrichment in featured deemed significant after differential expression analyses. Gene set enrichment analysis (GSEA) and over-representation analysis (ORA) used the “gseKEGG” and “enrichKEGG” functions from the “clusterProfiler” R package (version 4.8.3) ^76^. Gene Ontology (“BP”) enrichment analysis was conducted using “enrichGO” function from the “clusterProfiler” R package (version 4.8.3) and “org.Hs.eg.db” R package (version 3.17.0). The adjusted p-value was calculated using the Benjamini-Hochberg procedure ^74^ for multiple testing correction, with a significance threshold of 0.05 to identify enriched pathways.

### TIGER and PTM-SEA analysis

TIGER works by factoring the gene expression data in a population—represented as a genes-by-samples matrix—into a bipartite genes-by-transcription factor (TF) “gene regulatory network” (seeded with a prior network linking TFs and genes) and a TF by sample “transcription factor activity” matrix. In our analysis, we used the DoRothEA database ^77^ as the TF-gene binding prior knowledge, and we focused on TFs with high confidence levels as classified by the DoRothEA database, excluding TFs supported by only a single computational resource. We employed the TIGER function from the netZooR package (version 1.5.4) ^78^ for TF activity estimation. We employed the ‘TIGER’ function from the netZooR package (version 1.5.4) ^78^ for TF activity estimation. The estimation of kinase activity parallels that of TF activity; the primary difference involves replacing the expression matrix with a phosphorylation matrix and substituting the TF-gene binding prior with a kinase-substrate binding prior from the OmniPath R package (version 3.8.0) ^24^.

PTM-SEA, which is generally similar to GSEA, is a pathway enrichment method that detects activated or deactivated regulators like kinases or phosphatases through the enrichment of experimentally validated substrates ^34^. This tool leverages the PTMsigDB curated database ^34^, which offers the significant advantage of site-specific annotation and directionality of PTM regulation. We conducted PTM-SEA using the “ssGSEA2” R package (version 1.0.0) available on GitHub (https://github.com/broadinstitute/ssGSEA2.0), along with the PTMsigDB v2.0.0 database (https://proteomics.broadapps.org/ptmsigdb).

### Survival analysis

After performing GO enrichment analysis on sex-biased signaling proteins using CPTAC protein abundance data, we calculated a GO term score as the average protein abundance for each significantly enriched GO term. We used the R package “survival” (version 3.5.5) to fit Cox proportional hazards models to the CPTAC patients’ survival data. The resulting models were used to assess the impact of upstream signaling protein abundance, as quantified by the GO term score, along with sex effects and their interaction on survival outcomes, while adjusting for age, race, smoking status, tumor stage, and therapy. We used the “ggadjustedcurves” function with “average” method in the “survminer” R package (version 0.4.9) to make the adjusted survival curves. We also repeated this analysis on the target genes of TIGER gene regulatory network using TCGA gene expression data and TCGA patients’ survival data.

### Histone acetylation analysis

For the histone acetylation analysis, we included 67 detected acetylation sites: 32 on H1, 15 on H2A, 4 on H2B, 16 on H3, and none on H4, based on preprocessed acetylation data. We also included 15 detectable HATs and 13 HDACs from preprocessed protein abundance data. Lasso regression was applied to develop a regulatory network between histone regulators (HATs and HDACs) and histone acetylation sites.

Lasso regression is a linear regression technique that adds a regularization parameter (λ) multiplied by the sum of the absolute values of the model coefficients (the L1 norm, also known as the Manhattan distance) to the ordinary least squares loss function ^79^. This promotes sparsity by shrinking less important coefficients to zero, effectively performing feature selection. Each histone acetylation site was treated as the dependent variable, with HAT and HDAC proteins as covariates. Optimal λ values for each acetylation site were identified via 10-fold cross-validation to minimize mean squared error (MSE). To improve variable selection accuracy, this procedure was repeated 100 times using bootstrapping. Regression coefficients were selected when the 95th percentile of bootstrap coefficients exceeded 0.2 or the 5th percentile was less than -0.2. All analyses were performed using the R package glmnet (version 4.1.8).

## Code availability

R codes for all downstream analysis are available on a GitHub public repository: https://github.com/cchen22/Sex_Difference_LUAD.

## Acknowledgements

The results shown here are based upon data generated by the TCGA Research Network: https://www.cancer.gov/tcga. This work was supported by grants from the National Institutes of Health: CC, ES, JF, MBG, VF, KHS, and JQ were supported by R35CA220523; MBG and JQ were also supported by U24CA231846; JQ received additional support from P50CA127003; JQ and DLD were supported by R01HG011393; KHS and DLD were supported by P01HL114501; KHS was supported by T32HL007427; DLD was supported by K24HL171900; CMLR was supported by K01HL166376; CMLR and ES were also supported by the American Lung Association grant LCD-821824. MP was supported by R01CA251729. KG was supported by R01HL155749.

## Author contributions

CC: Conceptualization, Data curation, Formal analysis, Investigation, Methodology, Software, Validation, Visualization, Writing – original draft; ES: Data curation, Writing – review & editing; JF, MBG, VF, KHS, MP, KG: Writing – review & editing; DLD: Conceptualization, Writing – review & editing; CMLR: Conceptualization, Supervision, Writing – review & editing; JQ: Conceptualization, Funding acquisition, Resources, Supervision, Writing – review & editing.

## Supplementary Figures

Figure S1. Representative KEGG pathways enriched in the protein signaling network.

Figure S2. Survival analysis results using upstream signaling proteins and CPTAC.

Figure S3. Survival analysis results using downstream target genes and TCGA.

Figure S4. Sex-biased node degrees of HDACs in the regulation of histone acetylation.

Figure S5. Sex-biased drug effects of seven extra glucocorticoid receptor (NR3C1) agonists.

Figure S6. Sex-biased drug effects targeting sex hormone receptors – ESR1 and AR.

Figure S7. Sex-biased drug effects of MAPK inhibitors.

Figure S8. Adjusted p-values of limma differential expression analysis on key nodes.

Figure S9. Consistency check between validation datasets and discovery datasets.

Figure S10. Reproducibility of key nodes (AURKA, AR, NR3C1) using independent datasets.

## Supplementary Tables

Table S1. Sex by demographic variables cross table

Table S2. GSEA-KEGG pathway enrichment in differential gene expression: Female vs. Male

Table S3. GSEA-KEGG pathway enrichment in differential protein abundance: Female vs. Male

Table S4. ORA-KEGG pathway enrichment in protein phosphorylation: Female

Table S5. ORA-KEGG pathway enrichment in protein phosphorylation: Male

Table S6. ORA-KEGG pathway enrichment in protein acetylation: Female

Table S7. ORA-KEGG pathway enrichment in protein acetylation: Male

Table S8. ORA-KEGG pathway enrichment in the protein signaling subnetwork.

Table S9. ORA-GO pathway enrichment in the protein signaling subnetwork.

Table S10. ORA-GO pathway enrichment in the top 200 target genes.

